# GradeBins: quality control of bins from synthetic and real metagenomes

**DOI:** 10.64898/2026.03.11.709163

**Authors:** Brian Bushnell, Robert M. Bowers, Juan C. Villada

## Abstract

Genomes reconstructed from metagenomes support studies of microbial diversity and function, but their interpretation depends on how much DNA is recovered and whether foreign sequence is included. Quality can be measured in benchmarks, where contig origins are known, but must be estimated in routine analyses. Here we present GradeBins, which assesses individual bins (contigs assigned to one candidate genome) and summarizes the complete bin set from an assembly. For 6,374 simulated bins, its measurements agreed with AMBER and independent calculations within rounding. In control bins built from 20 kb fragments of reference genomes, GradeBins had lower observed completeness error than CheckM1 and CheckM2 and the highest observed F1 (the harmonic mean of precision and recall) for bin selection. With 2 kb fragments, GradeBins overestimated the completeness of incomplete bins, and its error increased from 3.1 to 17.6 percentage points. Contamination estimates differed among GradeBins, CheckM1, CheckM2 and GUNC, which define contamination differently. In sequenced metagenomes, the choice of estimator changed which bins passed quality cutoffs. GradeBins estimated quality faster than CheckM1, CheckM2 and GUNC on the same hardware. GradeBins thus provides fast quality assessment of prokaryotic bins and bin sets, with completeness estimates that are most accurate for long contigs.

## Introduction

Interpreting genomes reconstructed from metagenomes requires assessing their completeness and contamination^1,2^. These genomes reveal the metabolic potential of uncultivated microorganisms^3^ and expand the diversity represented in genome collections^4^. Metagenome-assembled genomes (MAGs) now describe microbial communities across Earth’s biomes^5,6^ and human populations^7,8^. Their value depends on the quality of the recovered sequence. Missing DNA limits the functions that can be inferred, whereas sequence from other organisms can create a misleading account of a genome’s content^1,9^.

Genome reconstruction involves decisions that affect both recovery and contamination. Assemblers such as MEGAHIT^10^ and metaSPAdes^11^ reconstruct contigs from mixed reads. Differences in contig coverage across samples help distinguish source genomes^12^. Binners including CONCOCT^13^, MaxBin 2.0^14^, MetaBAT2^15^ and QuickBin^16^ use coverage, sequence composition or other genomic features to group contigs. Neural network methods such as VAMB^17^, AAMB^18^, SemiBin^19^, SemiBin2^20^ and COMEBin^21^ provide further ways to learn these relationships. DAS Tool^22^ combines alternative assignments, and metaWRAP^23^ combines and reassembles bins. Assessing the resulting genomes is essential to comparing these workflows and choosing bins for subsequent analysis.

CheckM1^24^ and CheckM2^25^ estimate prokaryotic genome completeness and contamination, whereas GUNC^26^ assesses taxonomic consistency among genes. Contig coverage across samples can provide additional evidence of contamination^27^. The minimum information about a metagenome-assembled genome (MIMAG) standard^2^ combines completeness, contamination and RNA gene evidence into reporting categories. These assessments measure related but different quantities, so the same thresholds can select different genomes depending on the tool.

Workflow evaluation also concerns the complete collection of bins produced from an assembly, which we call a bin set. Two bin sets can contain the same number of genomes above a quality cutoff yet differ in DNA recovery, contamination and taxonomic representation. Sequencing technology and binning strategy^28^, as well as community complexity and sequencing depth^29^, affect genome recovery. Simulated communities provide known contig origins, which allow completeness and contamination to be measured directly. The Critical Assessment of Metagenome Interpretation (CAMI) challenge^30^ and its second round^31^ established shared benchmarks; CAMISIM^32^ and RandomReadsMG^33^ generate simulated data, and AMBER^34^ evaluates bin assignments. Benchmarks therefore rely on measurements from known contig origins, whereas routine analyses rely on estimates for genomes of unknown content. These two forms of assessment are usually provided by separate tools that differ in their definitions and outputs.

Here we present GradeBins, which provides both forms of assessment and reports the results for individual bins and for complete bin sets. We evaluate it with synthetic data, comprising bins from simulated communities and control bins constructed from reference genomes, and with bins from sequenced (real) metagenomes. We first test measurements made from known contig origins. We then examine quality estimates under controlled fragmentation and contamination, compare bin selection in sequenced metagenomes, and measure run time and RAM use. The analyses establish what GradeBins reports, how its estimates behave in the tested conditions and how those estimates affect the selection of bins.

## Results

### Reports for bins and bin sets

GradeBins takes existing bins and reports sequence statistics, completeness, contamination and quality classes. It assesses bins by one of two routes. For routine analyses of prokaryotic bins, it estimates completeness and contamination from bin FASTA files, together with an assembly or a size table and optional coverage and gene annotations (Fig. 1a). When contig origins and reference lengths are known, it measures completeness and contamination directly (Fig. 1b). Both routes produce quality classes and scores (Fig. 1c), a report for individual bins (Fig. 1d) and a summary of the complete bin set (Fig. 1e).

**Figure 1.**
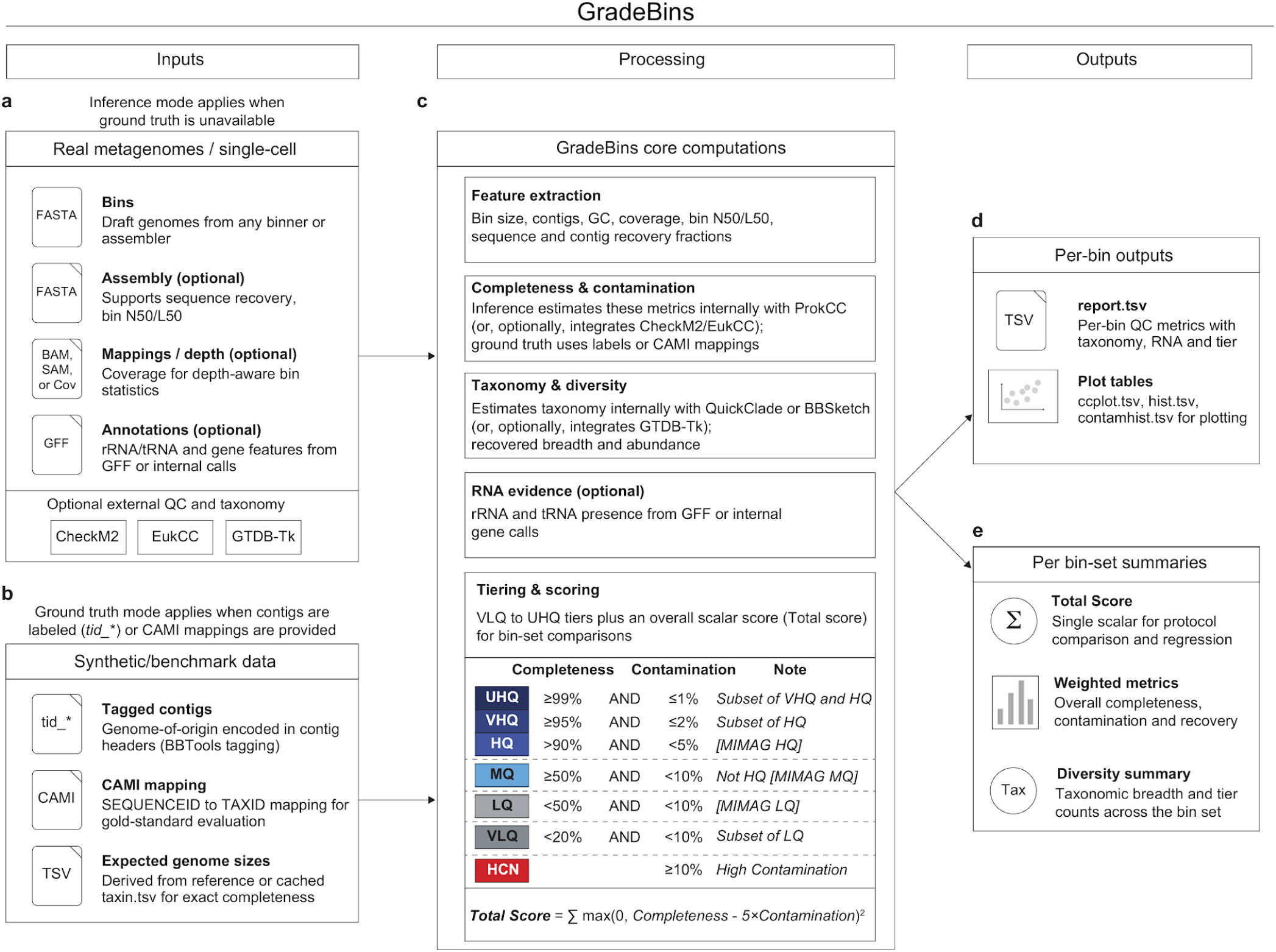
GradeBins inputs, calculations and outputs. **a,** Inputs for routine estimation (inference mode in the figure) are bin FASTA files and an assembly or size table, with optional coverage and gene annotations. The assembly is optional only when a size table supplies the required totals. Assemblies from individual cells are supported; this study evaluated metagenomic bins. **b,** Inputs for measurement from known contig origins (ground truth mode in the figure) are contig labels in FASTA headers or Critical Assessment of Metagenome Interpretation (CAMI) mappings, together with reference lengths. The validation used BBTools^35^ header labels; CAMI mappings are an alternative input. **c,** GradeBins calculates sequence statistics, completeness, contamination, quality classes (tiers in the figure) and scores. ProkCC supplies prokaryotic estimates; external quality and taxonomy reports are optional. UHQ, VHQ, HQ, MQ, LQ and VLQ denote ultra high, very high, high, medium, low and very low quality; HCN denotes high contamination. Colours identify classes. UHQ is within VHQ and HQ; VLQ is within LQ. Thresholds use completeness and contamination; full MIMAG high quality also requires RNA evidence. Total Score sums max(0, C − 5K)^2^ over bins, with C and K denoting completeness and contamination as fractions. **d,** The report describes individual bins and can supply plotting tables. Filenames identify output types; fields depend on inputs and enabled analyses. **e,** The summary describes recovery, quality counts, scores and taxonomic counts for the bin set. N/L statistics describe recovery across bins ordered by size. Methods define aggregate completeness and contamination scores. Taxonomic summaries count the taxa and bins represented. The evaluation of estimates used internal estimates for metagenomic bins.

Both routes start from the same bin FASTA files and lead to the same two outputs (Fig. 2a). Measurement refers to the main genome, the source genome that contributes the most DNA to a bin. For a bin containing 4 Mbp from a 5 Mbp main genome and 1 Mbp of foreign DNA, completeness is 80% and contamination is 20% (Fig. 2b). Estimation uses ProkCC (https://github.com/bbushnell/BBTools/blob/master/prokcc.sh; manuscript in preparation), the internal estimator of GradeBins, which predicts completeness and contamination from protein family counts and sequence features of each bin (Fig. 2c). The report records each bin’s size, contig count, GC content, quality estimates, class and gene counts (Fig. 2d). The summary describes assembly recovery, quality counts, bin sizes and scores (Fig. 2e). In the summary, some classes are nested within others. Ultra high quality is included in very high and high quality, and very low quality is included in low quality. The summary also gives two diagnostic counts for each class, Incomp and Contam, which count bins that meet class-specific conditions for incompleteness or contamination (Methods and Supplementary Methods); a bin can be counted in both.

**Figure 2.**
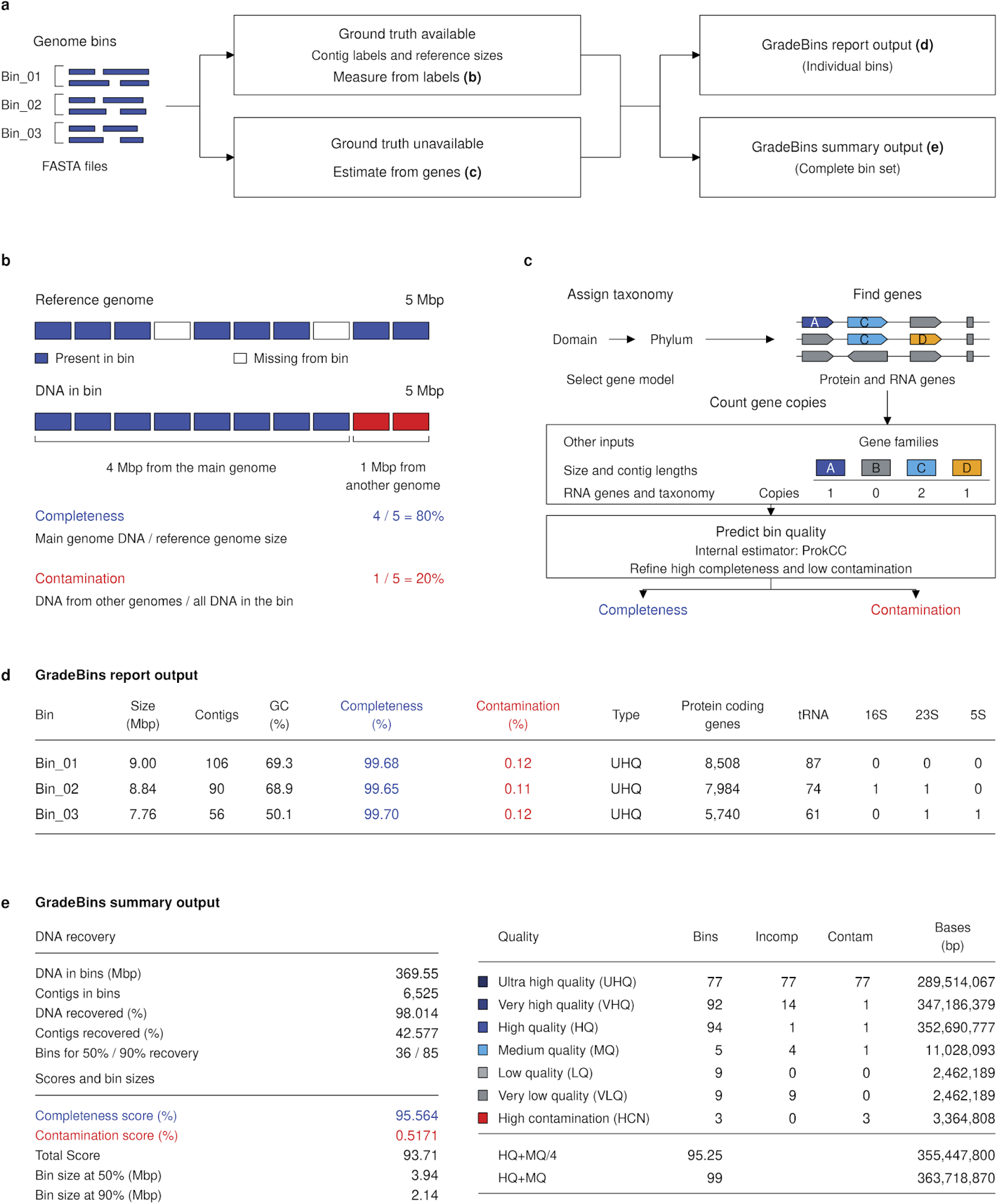
GradeBins assessment routes and example outputs. **a,** Bins are assessed by measurement from known contig origins (ground truth available) or by quality estimation (ground truth unavailable). Both routes produce a report for individual bins and a bin set summary. Bold letters refer to the example panels. **b,** Illustrative measurement. A bin contains 4 Mbp from a 5 Mbp main genome and 1 Mbp from another genome, giving 80% completeness and 20% contamination. Rectangles represent 0.5 Mbp; colours distinguish origins. **c,** GradeBins internal estimation with ProkCC. Taxonomy is assigned, a gene model is selected, genes are found and gene family copies are counted. ProkCC predicts completeness and contamination from these counts together with bin size, contig lengths, RNA genes and taxonomy, and then refines estimates of high completeness and low contamination. Matching colours connect illustrative genes to family counts. The drawing is a schematic of the procedure. **d,** Example report excerpt for the three largest bins in the MetaBAT2 bin set from the simulated community of 100 genomes. Bin_01–Bin_03 are display names. Completeness and contamination are estimates, Type is the quality class and gene columns give counts. **e,** Example summary for all bins in a bin set (*n* = 111). With bins ordered by size, the 50% and 90% statistics give the number of bins needed to recover that fraction of assembly DNA and the length of the last bin required. HQ+MQ sums counts; HQ+MQ/4 weights MQ by one quarter. UHQ is within VHQ and HQ; VLQ is within LQ. Blue shades identify HQ subdivisions; darker grey identifies VLQ. RNA criteria were disabled. Incomp and Contam are diagnostic counts within each exclusive class. All 77 UHQ bins are counted in both because they meet the UHQ cutoffs but have completeness <100% and contamination >0%. Blank cells are not applicable. Methods define thresholds and counting rules.

### Measurements agree with independent counts and AMBER

We tested the measurement route using five simulated prokaryotic communities with known contig origins (Supplementary Table 1); their planned source genomes are listed in Supplementary Table 2. Contigs were prepared by applying Tadpole from BBTools^35^ to each source genome separately. RandomReadsMG^33^ generated reads from the source genomes, and BBMap^36^ mapped them to the prepared contigs to supply coverage for binning. Two QuickBin^16^ configurations and one MetaBAT2^15^ configuration then binned each community, producing 15 bin sets that contained 6,374 bins (Fig. 3a). Because the contigs were prepared from individual genomes rather than from an assembly of mixed reads, every contig has a single known origin, and these inputs test how GradeBins evaluates bin assignments.

**Figure 3.**
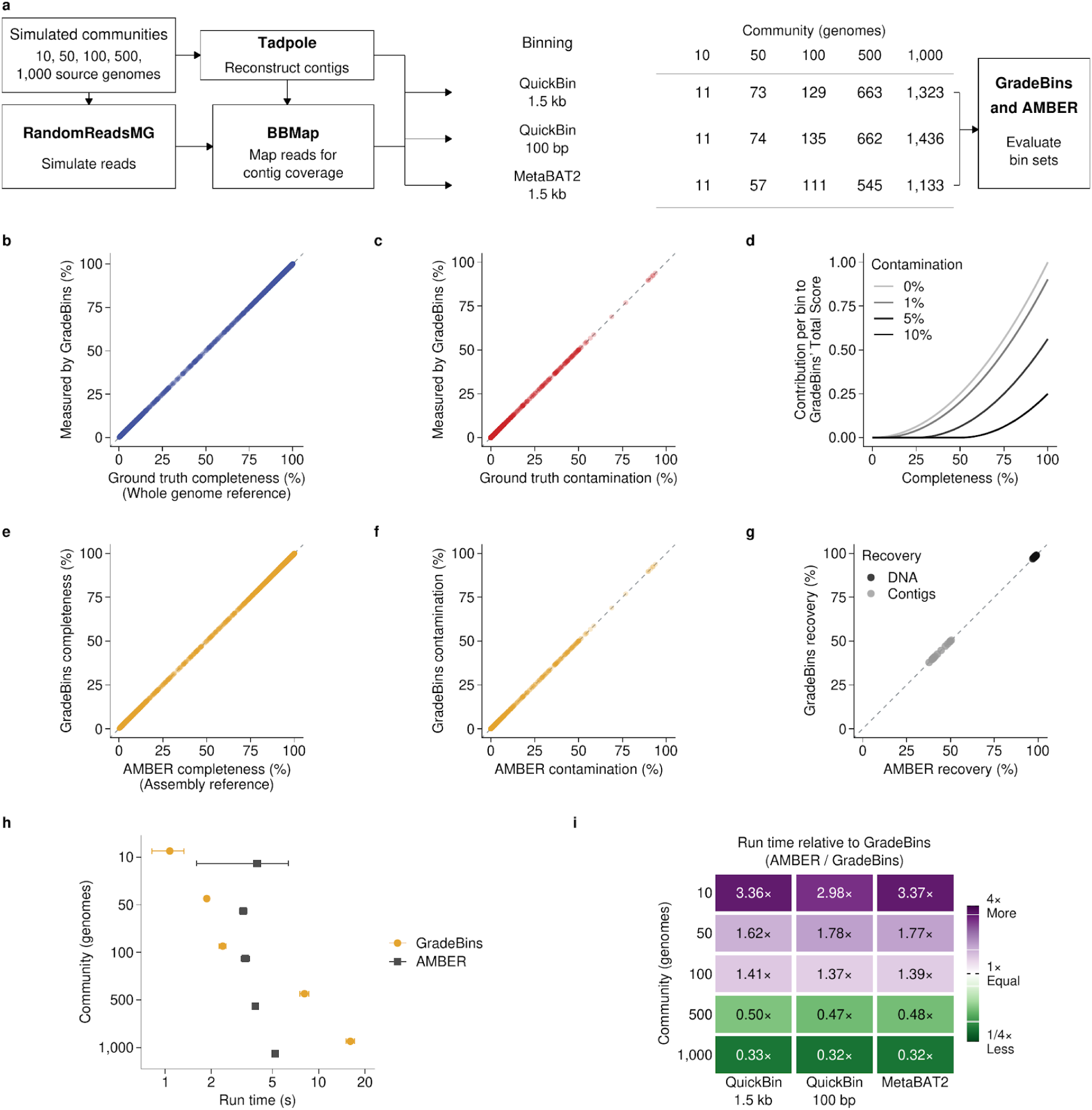
Measurements, scores and run time for bins with known contig origins. **a,** Five simulated prokaryotic communities were binned with two QuickBin configurations and MetaBAT2, giving 6,374 bins in 15 bin sets, each evaluated by GradeBins and AMBER. Labels give planned genome counts; the communities labelled 500 and 1,000 contain 497 and 998 represented genomes. **b,c,** Completeness and contamination measured by GradeBins against independent calculations. Completeness is DNA from the main genome (the largest contributor to a bin) divided by source genome length (whole genome reference); contamination is foreign DNA divided by total bin DNA. Sequence occurrences are counted, so completeness can exceed 100%. **d,** Each bin contributes max(0, C − 5K)^2^ to Total Score, with completeness C and contamination K as fractions. Darker grey indicates higher contamination. Contributions are summed without normalization or an upper limit. **e,f,** Completeness and contamination measured by GradeBins against AMBER. Both tools use the assembly reference, defined as all prepared contig DNA from the main genome, including contigs outside the bins. AMBER contamination is 100 × (1 − purity). Yellow identifies GradeBins. **g,** Percentages of assembly DNA (black) and contigs (grey) assigned to bins, from GradeBins and AMBER, for 15 bin sets. **h,** Run time of GradeBins (yellow circles) and AMBER (grey squares) by community, shown as means ± sample SD across nine runs per tool and community (three bin sets, each run three times). The time axis is logarithmic. **i,** AMBER run time divided by GradeBins run time for the same bin set and repetition; cells show medians of three ratios. On the logarithmic colour scale, white marks equality, green a shorter AMBER run time and purple a shorter GradeBins run time. Each dot in **b,c,e,f** is one bin (n = 6,374); diagonals in **b,c,e,f,g** mark equality. Run time includes startup, input loading and output generation. SD includes variation among bin sets and repetitions.

GradeBins measurements agreed with values recalculated by an independent parser, both for completeness, calculated as main genome DNA divided by source genome length (the whole genome reference), and for contamination, calculated as foreign DNA divided by total bin DNA (Fig. 3b,c). Agreement also held when completeness was calculated against the prepared contigs of the source genome (the assembly reference). For both references, the largest absolute difference was below 0.000503 percentage points, consistent with output rounding. Independent checks of sequence statistics and bin set summaries also agreed within their stated tolerances (Methods and Supplementary Methods). This validation establishes that GradeBins calculates these quantities correctly from the supplied labels and reference lengths.

GradeBins also reports a continuous Total Score for comparing bin sets (Fig. 3d). Each bin contributes max(0, C − 5K)^2^, where C and K are completeness and contamination as fractions. A complete bin without contamination contributes 1; ten such bins contribute 10. The score can therefore differ between bin sets that have identical quality counts, and it applies a chosen penalty for contamination.

We then compared GradeBins with AMBER^34^, a tool that evaluates bins with known contig origins, using the same 6,374 bins. With the assembly reference for completeness, the two tools differed by less than 0.00051 percentage points for completeness and for contamination in every bin, consistent with the rounding of GradeBins output (Fig. 3e,f). Both tools also agreed with values calculated independently from contig origins and integer base counts. For all 15 bin sets, the percentages of assembly DNA and of contigs assigned to bins agreed within the same tolerance (Fig. 3g). GradeBins and AMBER therefore give the same measurements of bin quality and recovery when contig origins are known.

Because the two tools gave the same measurements, we compared their run times on the same bin sets. Each tool was timed on its own input, bin FASTA files for GradeBins and tables of contig assignments for AMBER, and every run completed in under 18 seconds. GradeBins had the shorter mean run time for the communities of 10, 50 and 100 genomes (1.1 to 2.4 seconds, compared with 3.2 to 4.0 seconds for AMBER), whereas AMBER had the shorter mean run time for the communities labelled 500 and 1,000 (3.9 and 5.2 seconds, compared with 8.1 and 16.1 seconds for GradeBins; Fig. 3h). In all three binning configurations, AMBER took 1.37 to 3.37 times as long as GradeBins for the three smaller communities and 0.32 to 0.50 times as long for the two larger ones (median ratios; Fig. 3i). The relative run time of the two tools therefore depended on community size.

The two references used above give completeness different meanings. For example, a bin that recovers all prepared contigs of a source is 100% complete against the assembly reference but can be less complete against the whole genome reference (Extended Data Fig. 1a). Contamination is the same for both references because its denominator is total bin DNA. The tested bins covered a wide range of quality, as shown by their completeness against the assembly reference and their purity, the fraction of bin DNA from the main genome; the harmonic mean of these two quantities gives the F1 values in Extended Data Fig. 1b. These measurements require known contig origins. The next section examines the accuracy of estimates, which are used when contig origins are unknown.

**Extended Data Figure 1.**
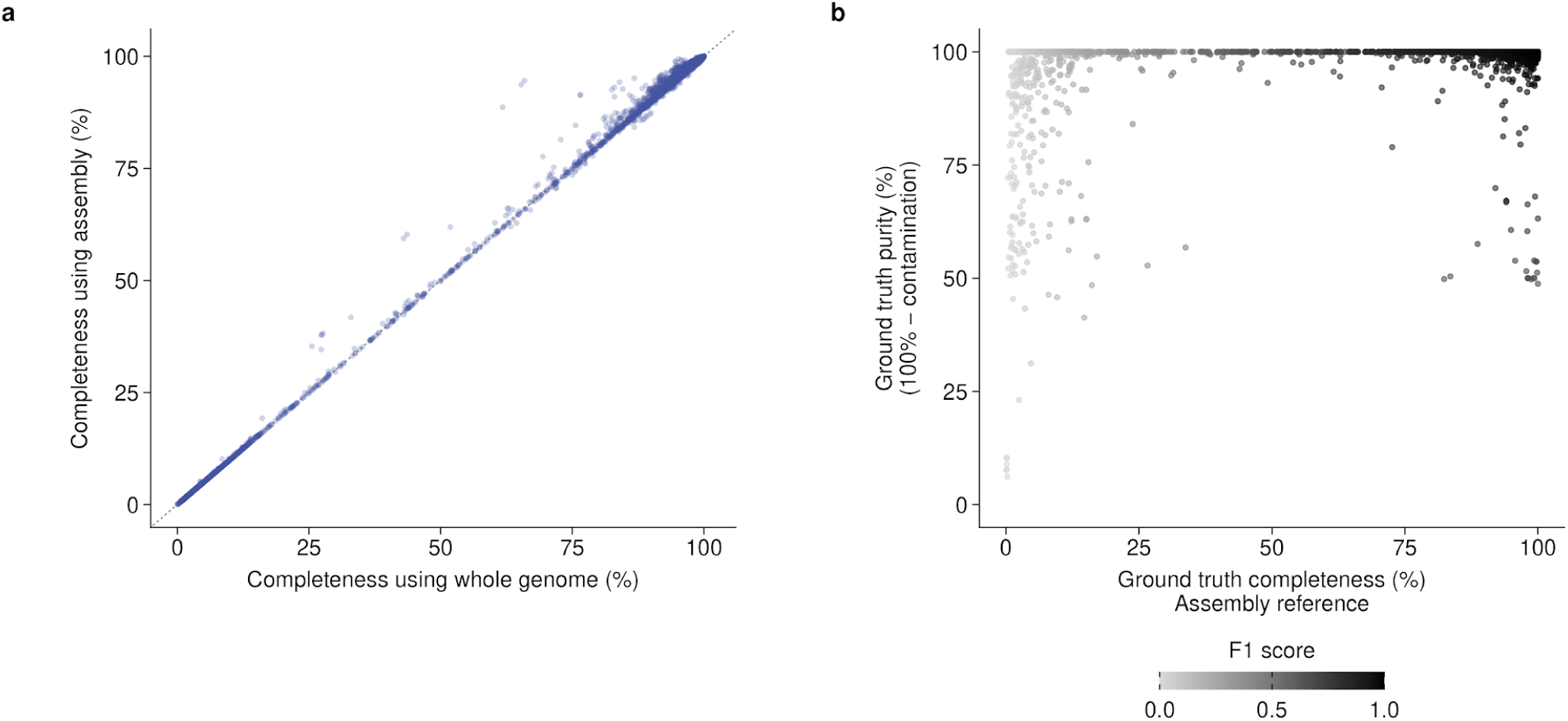
Completeness references and ground truth bin quality. **a,** Completeness for 6,374 bins against the whole genome reference (source genome length; horizontal axis) or the assembly reference (length of the prepared contigs of that source; vertical axis). Each point is one bin; the diagonal marks equality. Both calculations count sequence occurrences from the main genome. The prepared contigs can represent only part of the source genome. Figure 3b uses the whole genome reference, and Figure 3e uses the assembly reference. Contamination is the same for both references because its denominator is bin size. **b,** Completeness against the assembly reference and purity for the same bins. Purity is main genome DNA divided by bin size, equivalent to 100% minus contamination. These quantities are the recall and precision of the assignment of available sequence. Darker grey denotes higher F1, their harmonic mean. F1 was calculated independently from known contig origins for this analysis. The panel describes the quality of the tested bins.

### Fragment length affects accuracy

We next evaluated the accuracy of GradeBins estimates against ground truth that was defined independently of the tools. Because estimates describe genome content, ground truth for these tests counts each genome position once. For the simulated bins, DNA completeness is the fraction of main genome positions covered by the bin (Fig. 4a). We also constructed control bins (controls) from 26 reference genomes by cutting each genome into fragments of about 2 kb or 20 kb, retaining a set fraction of the fragments and adding a set fraction of foreign DNA; each fragment remained a separate sequence (Methods). For the controls, we compared the estimates with two ground truths. DNA completeness is the fraction of bases retained, and whole gene completeness is the fraction of annotated protein coding genes retained entirely (Fig. 4b,c). A gene split between fragments retains DNA but does not count as a whole gene. With 20 kb fragments, GradeBins estimates followed both ground truths. With 2 kb fragments, estimates for incomplete controls were higher than DNA completeness and exceeded whole gene completeness by a wider margin.

**Figure 4.**
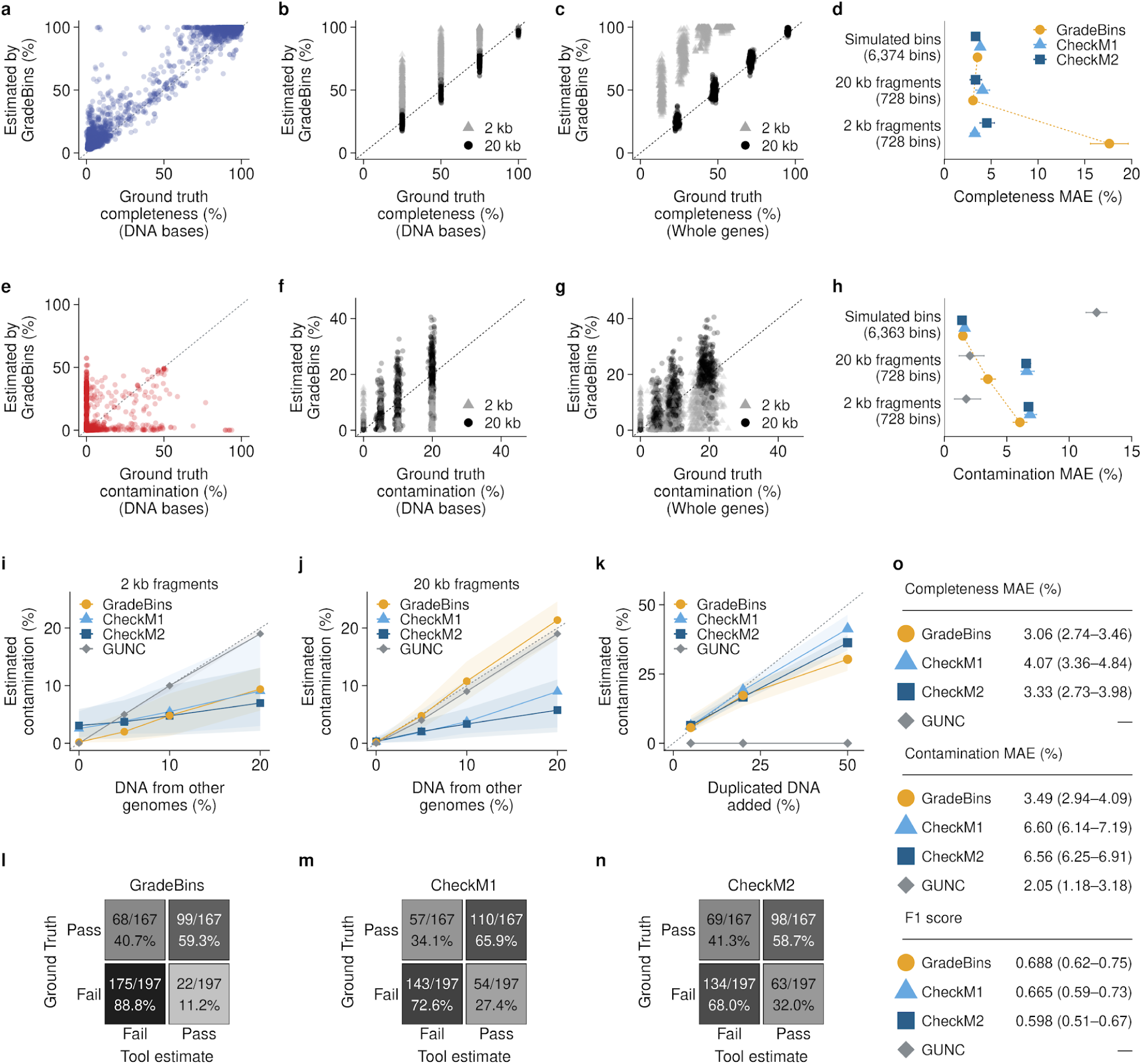
Quality estimates and bin selection in synthetic data. **a,** Completeness estimated by GradeBins against DNA ground truth for 6,374 simulated bins. **b,c,** Estimated completeness against retained DNA (**b**) or whole genes (**c**) for 2,912 controls (26 genomes, two seeds). Grey triangles, 2 kb fragments; black circles, 20 kb. **d,** Completeness mean absolute error (MAE) for simulated bins and 728 controls per fragment length (seed 4004). **e–g,** Estimated contamination against foreign DNA/total DNA (**e,f**) or whole foreign genes/all whole genes (**g**), using bins in **a–c**. Points are bins; diagonals mark equality. Blue, completeness; red, contamination. **h,** Contamination MAE for 6,363 simulated bins and the controls in **d**. **d,h** represent 1,552 simulated source genomes. **i,j,** Estimated contamination against requested foreign DNA with 2 kb (**i**) or 20 kb (**j**) fragments; 104 controls at zero, 208 per nonzero level. **k,** Estimated contamination against duplicated main genome DNA, as a percentage of the original bin; 52 controls per addition, none with foreign DNA. Lines and bands in **i–k** show medians and interquartile ranges. **l–n,** Bin selection for 364 additional 20 kb controls by GradeBins (**l**), CheckM1 (**m**), CheckM2 (**n**). Passing requires completeness >90% and contamination <5%, without RNA criteria. Rows give DNA ground truth (167 passing, 197 failing); cells show counts/row totals and row percentages. **o,** Summary for 20 kb controls, showing MAEs (**d,h**) and selection F1 (**l–n**). F1 is a fraction. Dashes mark unavailable quantities; GUNC estimates contamination only. Yellow circles, GradeBins; light blue triangles, CheckM1; dark blue squares, CheckM2; grey diamonds, GUNC. Yellow dashed lines connect GradeBins means. MAE (%) is in percentage points against DNA ground truth, with source genomes weighted equally; F1 pools bins. Error bars and parentheses show 95% percentile intervals from 2,000 source genome bootstrap samples. Contamination definitions differ among tools (Methods).

We summarized accuracy as the mean absolute error (MAE) against DNA ground truth and compared GradeBins with CheckM1 and CheckM2 on the same bins (Fig. 4d). In the simulated bins, completeness MAE was 3.53 percentage points for GradeBins, 3.82 for CheckM1 and 3.33 for CheckM2. In the controls, the ranking of the tools depended on fragment length. With 20 kb fragments, GradeBins had the lowest observed MAE, 3.06 percentage points (95% bootstrap interval 2.74–3.46), compared with 4.07 for CheckM1 and 3.33 for CheckM2. With 2 kb fragments, GradeBins MAE increased to 17.62 percentage points (15.60–19.64), whereas CheckM1 and CheckM2 had MAEs of 3.25 and 4.54. GradeBins completeness estimates were therefore close to ground truth for controls made of 20 kb fragments and too high for incomplete controls made of 2 kb fragments. MAE gives each source genome equal weight, and the intervals describe variation among the selected reference genomes.

We assessed contamination in the same simulated bins and controls (Fig. 4e–g). Ground truth is foreign DNA as a fraction of total bin DNA, or whole foreign genes as a fraction of all whole genes. Comparisons among tools used the DNA ground truth and the same bins for every tool, and here included GUNC (Fig. 4h). In the simulated bins, contamination MAE was 1.48 percentage points for GradeBins, 1.62 for CheckM1 and 1.41 for CheckM2, compared with 12.20 for GUNC. In the controls with 20 kb fragments, GradeBins MAE (3.49) was about half that of CheckM1 (6.60) and CheckM2 (6.56). GUNC had the lowest observed MAE in the controls at both fragment lengths (2.05 at 20 kb and 1.75 at 2 kb). These values measure agreement with DNA ground truth. Each tool keeps its own definition of contamination, and the definitions differ in the sequence features and denominators they use (Methods), so a difference from DNA ground truth can reflect the definition as well as estimation error.

We then examined how estimates responded to the amount of foreign DNA added to the controls. Median GradeBins estimates followed the added foreign DNA less closely with 2 kb fragments (Fig. 4i) than with 20 kb fragments (Fig. 4j). Adding duplicate copies of DNA from the main genome increased the median estimates of GradeBins, CheckM1 and CheckM2, although all DNA in these controls came from the main genome (Fig. 4k). Most GUNC estimates remained zero at every duplication level, which is consistent with its assessment of taxonomic mixture among genes. GradeBins and both CheckM versions therefore respond to redundant sequence from the same genome as well as to foreign sequence.

To test the consequences for bin selection, we applied the cutoffs completeness >90% and contamination <5% to 364 additional controls made from 20 kb fragments and designed to lie near these cutoffs. By DNA ground truth, 167 controls passed and 197 failed. GradeBins passed 99 of the 167 passing controls and 22 of the 197 failing controls; the corresponding counts were 110 and 54 for CheckM1 and 98 and 63 for CheckM2 (Fig. 4l–n). In this test, GradeBins therefore passed the fewest failing controls, whereas CheckM1 passed the most passing controls. Selection F1 was 0.688 for GradeBins (95% bootstrap interval 0.618–0.751), 0.665 for CheckM1 (0.587–0.730) and 0.598 for CheckM2 (0.512–0.670; Fig. 4o). The intervals overlap. F1 depends on the designed proportions of passing and failing controls. GUNC estimates contamination only, so Fig. 4o reports its contamination MAE.

### Estimator choice changes bin selection

We then compared GradeBins estimates with those of the other tools for bins from sequenced metagenomes. This analysis measures agreement among tools. It included 173 bins from five alternative bin sets of the MBARC-26 mock community^37^, distributed with the BinSPreader^38^ protocol^39^ in dataset version 3^40^, and 46 MAGs from one soil metagenome in the GEM catalogue^5^. Two features of these collections define the scope of the comparison. First, the MBARC-26 bin sets are alternative binnings of one assembly, so their bins are related. Second, the deposited soil MAGs had already passed CheckM filters for completeness and contamination. Each comparison pairs the estimates of two tools for the same bin, with separate plots and fitted lines for each tool and dataset. In soil, GradeBins completeness estimates covered a wider range than those of either CheckM version and were lower for some MAGs (Fig. 5a,b).

**Figure 5.**
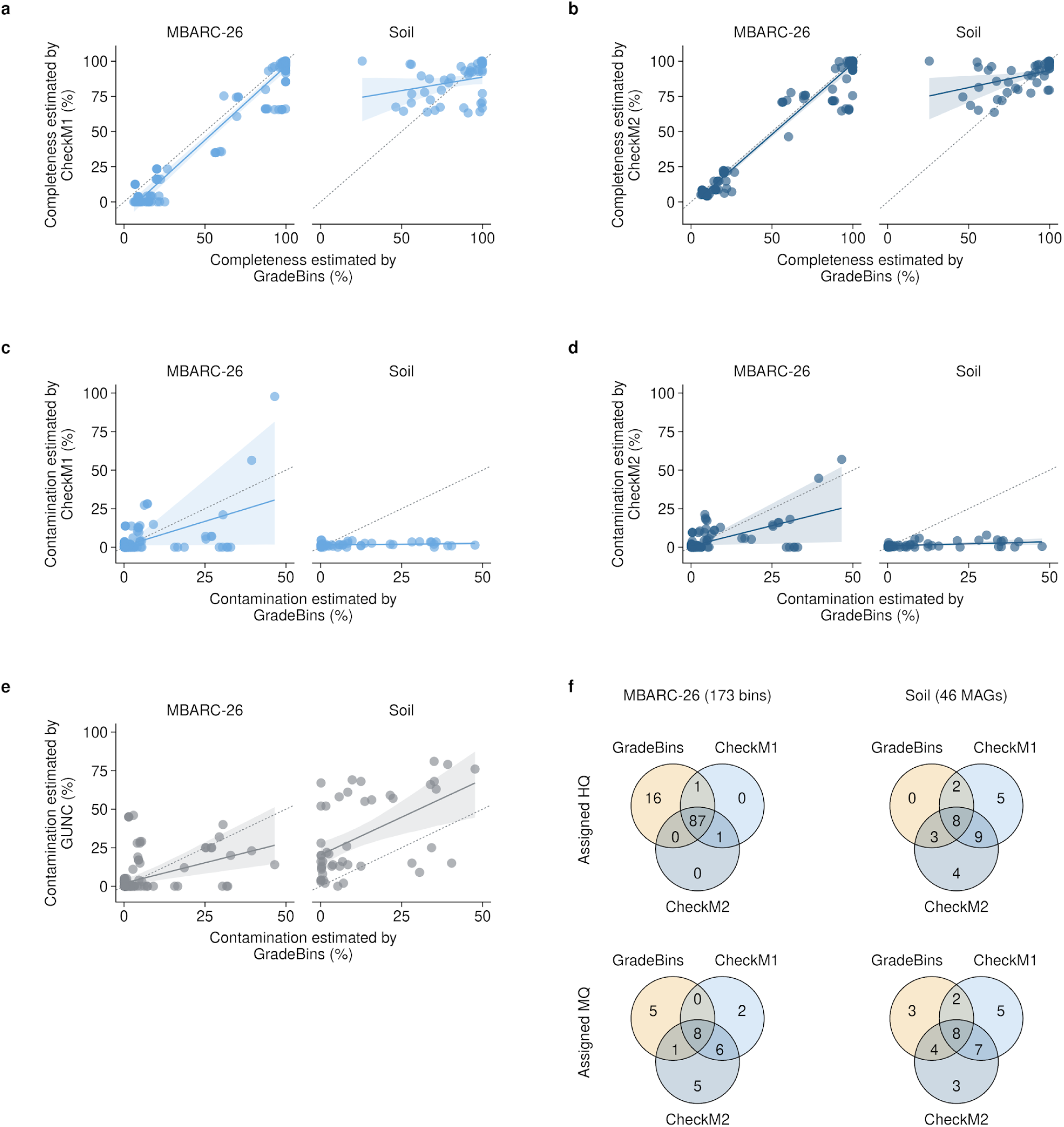
Quality estimates and assignments in sequenced metagenomes. **a,b,** GradeBins completeness compared with CheckM1 (**a**) and CheckM2 (**b**). **c–e,** GradeBins contamination compared with CheckM1 (**c**), CheckM2 (**d**) and GUNC (**e**). Each panel separates 173 MBARC-26 bins, from five alternative bin sets of one assembly, from 46 soil MAGs. Each dot pairs estimates for one bin. Light blue denotes CheckM1; dark blue, CheckM2; grey, GUNC. Dashed diagonals mark equal percentages. Contamination definitions differ among tools (Methods). Solid lines are separate ordinary least squares fits with intercepts for each tool, quantity and dataset. Shading shows pointwise 95% intervals for fitted means from 2,000 bootstrap samples of 35 groups defined by shared contigs for MBARC-26 and of 46 MAGs for soil. Lines and bands span observed GradeBins values. Bands describe fitted means, rather than individual predictions. **f,** HQ (top) and MQ (bottom) assignments for MBARC-26 (left) and soil (right). Yellow denotes GradeBins; CheckM colours match **a–d**. Numbers count bins in each exact combination, including zeros. Circle areas are schematic. Each diagram contains the bins assigned to the class by at least one tool. HQ requires completeness >90% and contamination <5%; MQ requires completeness ≥50% and contamination <10%, excluding each tool’s HQ bins. Assignments use original estimates. Full MIMAG high quality additionally requires RNA evidence, which was not assessed. GUNC provides no completeness estimate or combined assignment. MAG, metagenome-assembled genome.

For most soil MAGs, GradeBins estimated more contamination than either CheckM version (Fig. 5c,d), and GUNC reported more contamination than GradeBins (Fig. 5e). Because these MAGs had been selected using CheckM estimates, the comparison is conditioned on low CheckM contamination. These comparisons describe agreement among estimates, including the effects of different contamination definitions. Identifying the estimate closest to genome content would require assigning the bins to reference genomes.

These differences changed quality assignments when the estimates of each tool were classified with the same completeness and contamination cutoffs from MIMAG^2^. For MBARC-26, GradeBins assigned 104 bins to high quality (HQ), compared with 89 for CheckM1 and 88 for CheckM2; 87 bins were HQ for all three tools. For soil, the direction was reversed, and GradeBins assigned 13 MAGs to HQ, compared with 24 for either CheckM version. GradeBins assigned 17 soil MAGs to medium quality (MQ), compared with 22 for either CheckM version, and its contamination estimates for the remaining 16 soil MAGs were at least 10%. MQ assignments also differed for MBARC-26 (Fig. 5f).

These counts show that the choice of estimator changes which bins are retained, and that the direction of the change differed between the two collections. The assignments use completeness and contamination only; full MIMAG high quality designation additionally requires RNA gene evidence. Because the MBARC-26 bin sets are related and the soil MAGs had been filtered with CheckM, the comparison describes how estimator choice affects selection within these two collections.

### GradeBins estimates bin quality faster across tested datasets

We first examined how GradeBins run time changed with input size, using the 45 runs that had assessed the 15 simulated bin sets in three modes. Measurement from known contig origins completed within one minute in every run, whereas quality estimation took several minutes for the largest inputs (Fig. 6a). These runs used different hardware and were each performed once, so they describe the recorded workloads.

**Figure 6.**
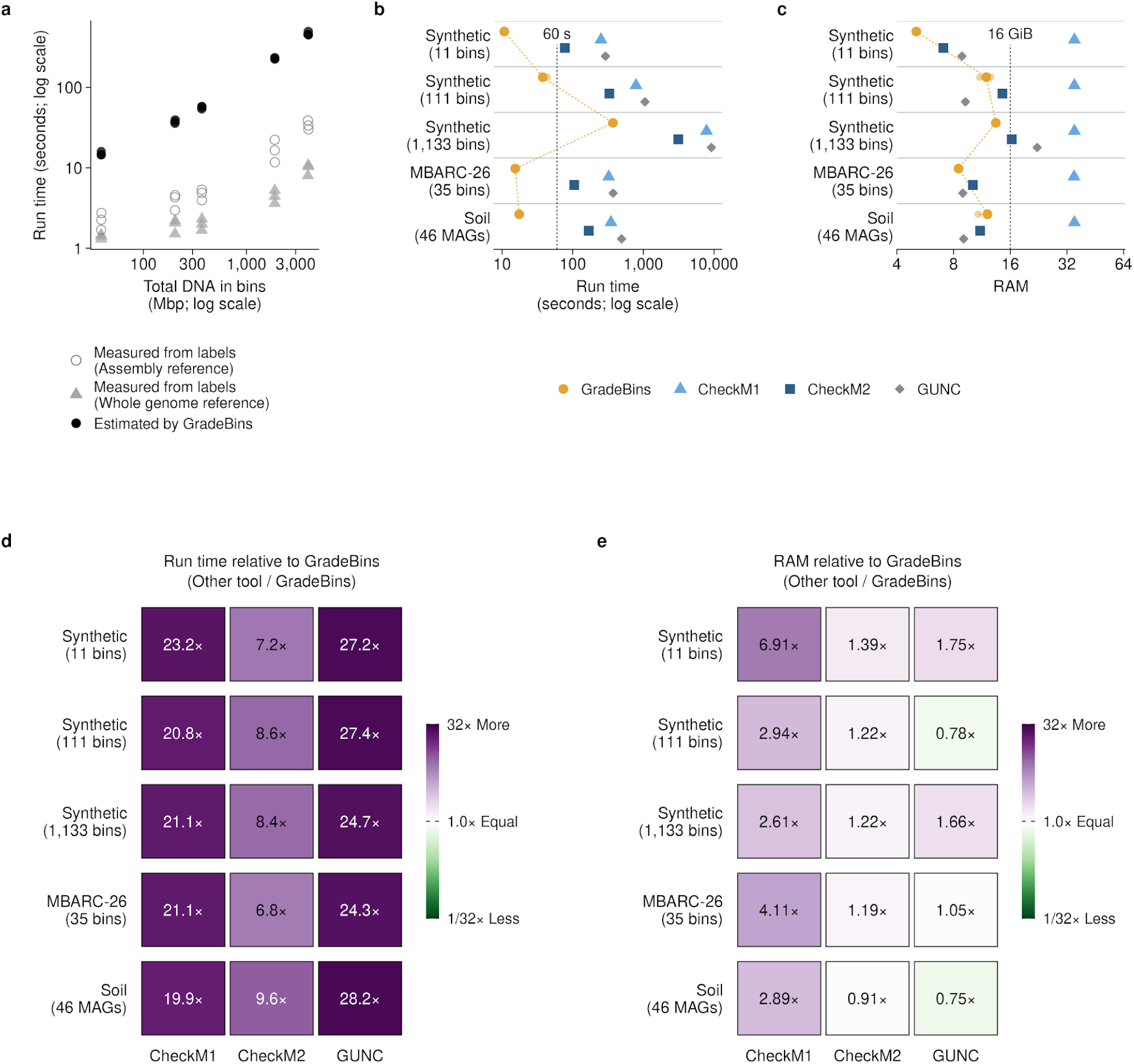
GradeBins run time and RAM use. **a,** GradeBins run time for 45 runs on the 15 simulated bin sets, 15 per mode. Open grey circles, measurement from contig labels with the assembly reference; light grey triangles, measurement from contig labels with the whole genome reference; black circles, quality estimation. Input size is total bin DNA. Hardware differed and each run was performed once. Axes are logarithmic. **b,** Run time for quality estimation on five datasets, which were simulated bin sets of 11, 111 and 1,133 bins (labelled Synthetic), 35 MBARC-26 bins and 46 soil MAGs. Within each dataset and repetition, all tools used the same node and 16 logical CPUs; the seven nodes used had the same processor model. Small dots show three runs, large symbols their medians and bars their ranges. All 60 runs completed. Yellow circles, GradeBins; light blue triangles, CheckM1; dark blue squares, CheckM2; grey diamonds, GUNC. Yellow dashes connect GradeBins medians. The vertical line marks 60 seconds. **c,** GNU time maximum resident set size (RSS), in GiB (2^30^ bytes), with symbols as in **b**. This records the largest individual process peak, including completed helpers. The dashed line marks the 16 GiB Java heap setting, distinct from total RAM. Axes in **b,c** are logarithmic. **d,e,** Run time (**d**) or RAM (**e**) of each comparison tool divided by that of GradeBins. Cells show medians of three ratios calculated within each dataset and repetition. White marks 1× on a logarithmic colour scale; green denotes that the comparison tool used less than GradeBins and purple that it used more. Run time includes startup and model loading and excludes database preparation. GradeBins run time includes waiting for its taxonomy service; RAM measurements cover local processes. Extended Data Figure 2 compares RSS with the combined RAM of the tool and its helpers.

For a controlled comparison of quality estimation, we ran GradeBins, CheckM1, CheckM2 and GUNC three times on each of five datasets. Within each dataset and repetition, all four tools ran on the same reserved compute node with 16 logical CPUs. Nodes differed among repetitions but had the same processor model. GradeBins had the shortest run time in all 15 comparisons. Its median run time ranged from 10.76 to 37.72 seconds for four datasets and was 371.27 seconds for the largest dataset of 1,133 bins (Fig. 6b). Its median peak RAM, measured as the maximum resident set size recorded by GNU time^41^, ranged from 5.07 to 13.41 GiB (Fig. 6c).

For each dataset and repetition, we divided the run time or peak RAM of each comparison tool by the corresponding GradeBins value. CheckM2 was the fastest comparison tool, and its median run time was 6.81 to 9.65 times that of GradeBins across datasets; the median ratios were 19.9 to 23.2 for CheckM1 and 24.3 to 28.2 for GUNC (Fig. 6d). GradeBins used less peak RAM than CheckM1 in every dataset, whereas it used more or less than CheckM2 and GUNC depending on the dataset (Fig. 6e).

GNU time records the peak of the largest single process. As a complementary measurement, we sampled the combined RAM of each tool and its helper processes (Extended Data Fig. 2a). The two measurements gave different values and ratios, because one is the peak of a single process and the other a sampled sum across processes (Extended Data Fig. 2b,c). Both cover local processes. GradeBins run time includes waiting for its remote taxonomy service, whose resources lie outside these measurements. The comparisons therefore describe local resource requirements under the tested conditions.

**Extended Data Figure 2.**
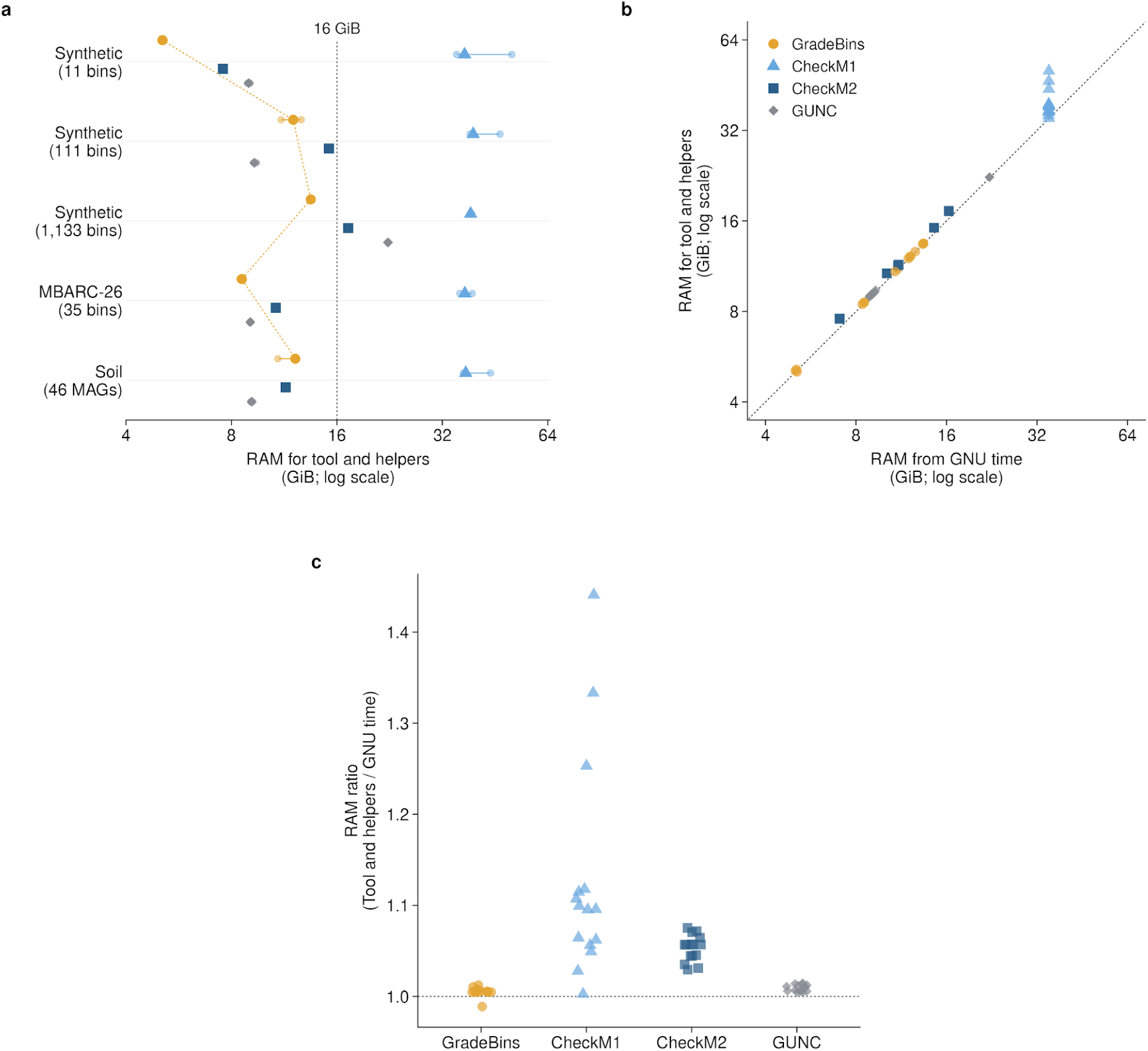
Combined memory use and GNU time measurements. **a,** Largest sampled combined proportional set size (PSS) for the tool and helpers in the same 60 runs as Figure 6b,c. PSS apportions shared memory among processes. Small dots show three runs, large symbols their medians and bars their ranges. Yellow dashed lines connect GradeBins medians. The vertical line marks the 16 GiB Java heap setting. One GiB is 2^30^ bytes. **b,** GNU time maximum resident set size (RSS) versus sampled combined PSS. Each point is one run (n = 60). Axes have identical ranges and logarithmic scales; the diagonal marks equality. GradeBins uses yellow circles; CheckM1, light blue triangles; CheckM2, dark blue squares; GUNC, grey diamonds. The key applies to **a–c**. RSS records an individual process peak; combined PSS sums across the tool and helpers, with different accounting and sampling. **c,** Combined PSS peak/GNU time maximum RSS, with 15 runs per tool across five datasets and three repetitions. The line at 1 marks equality. Peaks can occur at different times and ratios can be below 1. Measurements cover local processes. Sampling may miss brief peaks; Methods describe the procedure and observed intervals.

## Discussion

GradeBins reports measurements from known contig origins and estimates of prokaryotic bin quality in the same report and summary formats. Its measurements agreed with AMBER and independent counts when contig origins were known. In the simulated bins and in controls made of 20 kb fragments, the observed errors of its estimates were similar to or lower than those of CheckM1 and CheckM2, whereas it overestimated the completeness of incomplete controls made of 2 kb fragments. In sequenced metagenomes, the choice of estimator changed which bins were retained. GradeBins also estimated quality faster than CheckM1, CheckM2 and GUNC in the controlled comparisons, supporting repeated assessment during workflow development. For measurements from known contig origins, GradeBins was faster than AMBER for the three smaller communities, whereas AMBER was faster for the two larger ones.

The accuracy of an estimate depends on the quantity used as ground truth. Measurements from contig labels count sequence occurrences relative to the supplied reference lengths, whereas our tests of estimates counted unique genome coordinates and whole genes. These quantities can differ for the same bin. Recovery can also vary among sequence features; for example, short read binning can recover plasmids and genomic islands less well than chromosomes^42^. The design of the tests defines the scope of these results. Our simulated contigs were prepared separately from each source genome, which isolates the evaluation of bin assignments; assembly from mixed reads adds further sources of error, which are represented in CAMI II^31^ and other benchmarks^29^. Fragment length had a large effect on GradeBins completeness estimates, which were closest to ground truth for long fragments; contig length should therefore be considered when interpreting GradeBins estimates. Estimation used the current reference of a remote taxonomy service at the time of each run. Performance should be interpreted within these tested conditions.

Total Score provides a continuous summary for comparing bin sets from the same assembly. Its value combines the number and quality of bins under a fixed contamination penalty. Quality scores also guide methods that combine bins^22^ or remove redundant reconstructions^43^. The Total Score penalty and the separate HQ+MQ/4 weighting are chosen reporting weights, rather than empirically established measures of scientific value. They are best interpreted alongside sequence recovery, contamination and the requirements of the intended analysis. Likewise, the ultra high and very high quality classes of GradeBins apply additional thresholds that are 1 to 5 percentage points apart. The MAEs observed for GradeBins estimates under the most favourable conditions tested (about 1.5 to 3.5 percentage points) are of similar size. For estimated values, these classes therefore serve as descriptive labels.

Long read sequencing^44,45^ and assembly methods such as metaMDBG^46^ enable more continuous genome reconstructions and expand the diversity recovered across habitats^47^. Such assemblies have longer contigs, the condition under which GradeBins estimates were most accurate in our controls. Greater continuity can also improve recovery of RNA operons and other genomic elements^48^. RNA evidence remains useful even for genomes with high estimated completeness, because earlier catalogues reported many MAGs without a full rRNA complement^49^, which is required for MIMAG high quality designation^2^. GradeBins reports RNA gene counts alongside completeness and contamination. Applying the full standard also requires checking the specified rRNAs and tRNAs for at least 18 amino acids; a count of 18 tRNA genes alone does not establish that coverage.

Differences among quality estimates affect reference resources as well as individual studies. GTDB release 10^50^ reported changes in genomes passing quality filters after adopting CheckM2 and revising its eligibility rules. Our comparison in sequenced metagenomes shows the consequences of estimator choice within the tested collections; establishing the true genome content of these bins would require independent evidence. Accessory genes and horizontal transfer^1^, and gaps in reference databases^26^, complicate the interpretation of contamination estimates. Sequence inspection and reference comparisons^51^ provide complementary evidence. Genomes from individual cells can link gene content to organisms independently of metagenomic binning^52,53^, although GradeBins performance on those assemblies remains to be evaluated.

Quality assessment should report both the evidence for each bin and the outcome for the complete bin set, and GradeBins provides both. When bin sets are compared, the same reference lengths, ground truth definitions and settings should be used. The estimator and cutoffs used to select genomes should also be reported, because these choices affect subsequent analyses.

## Methods

### GradeBins execution

We evaluated GradeBins in BBTools 40.04^35^ using existing bin FASTA files. Internal prokaryotic quality estimation used ProkCC (https://github.com/bbushnell/BBTools/blob/master/prokcc.sh; manuscript in preparation), selected with prokcc, 16 threads and a 16 GiB maximum Java heap. Simulated bins and controls used anonymous contig identifiers; sequenced MBARC-26 and soil bins retained their public identifiers. Origin labels and comparison tool outputs were excluded from estimation inputs.

Version 40.04 requires an assembly (ref=) or size table (taxin=), including for estimation. Measured input sizes supplied these totals for controls and soil MAGs. We interpreted assembly recovery only for bin sets with a corresponding assembly, because each control is a separate test without a shared assembly and the original soil assembly was unavailable. Gene calling and quality estimation ran locally, with taxonomic context requested from the remote BBTools QuickClade^35^ service. The outer quickclade=f option disables a separate taxonomy step but leaves this internal request active. Run time includes waiting for the service; RAM measurements exclude remote resources. Each run used the current reference of the remote service.

BBTools ProkCC^35^ combined protein family counts and sequence features as inputs to local prediction models. GradeBins used the fraction-valued outputs named gene_completeness and gene_contamination; separate base pair outputs were not used. The DNA and whole gene ground truths used for evaluation are defined independently below.

Optional imports include quality reports from CheckM2^25^ and EukCC^54^ and taxonomy from GTDB-Tk^55,56^. We evaluated internal estimates for prokaryotic metagenomic bins; external report imports and assemblies from individual cells were outside this evaluation.

### Measurements from known contig origins

The main genome contributes the most bases to a bin. Let D be its bases in the bin, B total bin length, G source genome length and A prepared contig length from that source. Completeness is D/G for the whole genome reference or D/A for the assembly reference; contamination is (B − D)/B for either. Percentages are 100 times these fractions. Contig headers or CAMI mappings supply source identifiers. We validated the BBTools tid_* labels, which contain NCBI Taxonomy^57^ identifiers, using supplied reference lengths.

These measurements count sequence occurrences, including repeats, so completeness can exceed 100%. Recovering all prepared contigs gives D/A = 1 even when D/G < 1. Source FASTAs define G; draft sources represent supplied sequence rather than independently complete genomes. Figure 3b uses D/G, Figure 3e uses D/A, and Extended Data Figure 1a compares the two references.

Extended Data Figure 1b uses D/A as recall and D/B as precision (purity). F1 is twice their product divided by their sum, using fractions. This F1 was calculated from known contig origins for this analysis, separately from GradeBins report fields.

### Report fields, recovery and scores

Figure 2d,e shows the three largest bins and the complete set of 111 bins in the MetaBAT2 bin set from the simulated community of 100 genomes. Display names Bin_01–Bin_03 correspond to bin.41, bin.110 and bin.46. The excerpt retains the rounded values of the report and omits unavailable fields. Supplementary Methods define report fields and diagnostic counts.

DNA recovery (Sequence Recovery in GradeBins output) is 100 × total bin length/assembly length; contig recovery is 100 × binned contig count/assembly contig count. For bins ordered by decreasing length, each N/L recovery threshold reports the number of bins needed to reach that fraction of assembly DNA and the last required bin length. By default, GradeBins uses N for count and L for length; swapnl=t exchanges these labels.

For bin length L, completeness C and contamination K as fractions, contaminating bases F = round(K × L). With assembly length T, the completeness score is 100 × sum(round(C × (L − F)))/T and the contamination score is 100 × sum(F)/T. Both retain rounding to whole bases and the assembly denominator.

Total Score = sum(max(0, C − 5K)^2^). A complete uncontaminated bin contributes 1. The sum is unnormalized and uncapped; contributions can exceed 1 when completeness exceeds 1. The multiplier and exponent are reporting choices. Figure 3d illustrates C = 0–1 and K = 0, 0.01, 0.05 and 0.10.

### Quality classes and diagnostic counts

RNA criteria were disabled for the illustrated report and selection comparisons. Thresholds for completeness and contamination were ≥99% and ≤1% for UHQ; ≥95% and ≤2% for VHQ; >90% and <5% for HQ; ≥50% and <10% for MQ, excluding HQ; <50% and <10% for LQ; and <20% and <10% for VLQ. The threshold for HCN was contamination ≥10%. The abbreviations denote ultra high, very high, high, medium, low and very low quality, and high contamination. Full MIMAG high quality^2^ also requires 23S, 16S and 5S rRNA genes and tRNAs for at least 18 amino acids. The RNA criterion in GradeBins counts tRNA genes, whereas full MIMAG designation requires their amino acid identities.

Each bin receives one exclusive report Type. In the summary, the Bins and Bases columns are nested for UHQ, VHQ and HQ and for VLQ and LQ; HQ, MQ, LQ and HCN do not overlap.

Incomp and Contam count diagnostic conditions within exclusive report Types and can overlap (Supplementary Methods). All 77 UHQ bins in Figure 2e have completeness <100% and contamination >0% while satisfying UHQ thresholds.

HQ+MQ sums the bin counts; HQ+MQ/4 assigns MQ a chosen weight of one quarter. Its Bases value likewise uses HQ bases plus MQ bases divided by four, with integer division. Taxonomic summaries count represented taxa and bins, rather than organism abundance.

### Simulated communities and binning inputs

Five simulated prokaryotic communities had planned sizes of 10, 50, 100, 500 and 1,000 genomes; in the last two, 497 and 998 genomes were represented in the prepared contigs, because three and two planned genomes had no prepared contigs (Supplementary Table 1). Supplementary Table 2 lists the planned source genome members, taxonomy identifiers, assembled sequence lengths and whether each was represented in the prepared contigs. Reassemble applied Tadpole separately to each source FASTA with BBTools 39.70^35^ and k=155. Contigs were combined and sorted by decreasing length. Thus, preparation reconstructed individual source genomes rather than assembling mixed metagenomic reads.

RandomReadsMG^33^ generated four sets of paired 150 bp reads per community. Commands specified depth limits of 0.5–64, alternating min4 and exp abundance modes, sequence errors, sinusoidal coverage variation with waveamp=0.5 and origin bias of 0.1. BBMap^36^ mapped each set to the contigs with maxindel=3 and ambig=random.

Each community supplied two QuickBin^16^ runs and one MetaBAT2^15^ run. QuickBin used mincontig=1500 or the 100 bp default, threads=16 and mincluster=50000. The 100 bp default is documented in the preserved BBTools 39.76 script and DataLoader source; an exact BBTools release was not recorded for every historical QuickBin run. MetaBAT2 2.18 used minimum contig and cluster lengths of 1,500 and 200,000 bp. QuickBin received four BAM files; MetaBAT2 received their depth summary. Historical QuickBin runs also differed in software paths and validation settings, so their differences cannot be attributed solely to minimum contig length.

GradeBins 40.04 evaluated 6,374 bins from 15 bin sets. Each community provided three alternative bin sets of the same contigs. Reports that use the whole genome or the assembly reference describe the same bins.

### Validation of reported measurements

An independent FASTA parser counted bases by source label and recalculated completeness and contamination against both references, covering 12,748 reports for 6,374 bins. The maximum absolute discrepancy was 0.00050235 percentage points, consistent with printed precision. Values were not clipped at 100%.

We also checked size, contig count and GC content in 25,542 reports, including repeated reports for the simulated bins and reports for 46 soil MAGs. Counts had to agree exactly; GC fractions had a tolerance of 0.0005001. Assembly recovery checks used 15 bin sets, and N/L checks used 11 recovery thresholds per set. All 76,986 additional checks met their specified criteria (Supplementary Methods). Repeated reports are not additional biological samples.

### Comparison with AMBER using known contig origins

AMBER^34^ version 2.0.8 evaluated the same 6,374 bins in 15 bin sets from the five simulated prokaryotic communities. The three bin sets of each community are alternative binnings of the same prepared contigs. The comparison reused the existing reads, contigs, bin assignments and GradeBins 40.04 reports.

For each community, a CAMI format ground truth file recorded every prepared contig, its source genome and its length, including contigs absent from the bins. A separate file recorded the contigs assigned to each bin. Sequence identifiers were linked to the original FASTA headers through the existing contig table, and the length of each assigned contig was checked against its ground truth record. Each contig occurred in at most one bin within a bin set, and each bin had one source contributing the most bases.

The unmodified cami-amber 2.0.8 release ran in an isolated Python 3.11 environment with `amber.py -g ground_truth.tsv -l “Input bins” -p 0 -n 0 --skip_gs --stdout -o amber bins.tsv`. The settings retained all supplied assignments and contig lengths; --skip_gs omitted evaluation of the ground truth as a separate binning result. Numerical comparisons used the output tables written by AMBER.

Using the definitions above, AMBER completeness is D/A and purity is D/B. We expressed completeness as a percentage and calculated contamination as 100 × (1 − purity). The GradeBins reports using the assembly reference use the same completeness denominator and calculate contamination as 100 × (B − D)/B. Independent values were calculated from source labels and integer base counts. Thus, Figure 3e uses the assembly reference for both tools, whereas Figure 3b uses the whole genome reference.

For each bin set, AMBER reports the fractions of assembly bases and contigs assigned to bins. After conversion to percentages, these were compared with the GradeBins Sequence Recovery and Contig Recovery fields and with independently calculated assigned bases or contigs divided by their assembly totals. Because each contig was assigned to at most one bin, the recovery definitions of the two tools are equivalent for these inputs. The comparison used values for individual bins and bin sets. AMBER also reports average completeness across source genomes, which is averaged over genomes and is therefore a separate quantity from the values for individual bins.

We calculated absolute differences between the tools, and between each tool and the independently calculated values, for every bin and bin set. Printed values were compared with the independent values using tolerances of 0.00051 percentage points for GradeBins, which reflects its output rounding, and 10^−10^ percentage points for AMBER. All values were within these tolerances, and the largest difference between the tools was 0.000502 percentage points. These comparisons are descriptive and include all results. Source Data Figure 3 provides the reported measurements, integer counts, reference lengths and differences.

### Run time for measurements from known contig origins

For the comparison in Figure 3h,i, we ran GradeBins 40.04 and AMBER 2.0.8 three times for each of the 15 bin sets, giving 90 runs. All runs were sequential on one exclusively allocated node with an AMD EPYC 9555P processor, and both tools were restricted to the same 16 logical CPUs. GradeBins used 16 threads and a 16 GiB maximum Java heap. Numerical library thread settings were fixed at one for both tools. The 45 combinations of bin set and repetition were shuffled using seed 4040311, and tool order was shuffled within each combination.

GNU time^41^ measured complete runs, including startup, input loading and output generation. GradeBins read bin FASTA files and assembly reference information; AMBER read CAMI tables. Input preparation and conversion were excluded. Each run used a fresh output directory. Operating system caches were not flushed and no warm-up runs were performed. All 90 runs completed.

Figure 3h shows arithmetic means and sample SD across nine runs per tool and community (three bin sets, each run three times). The SD therefore includes variation among bin sets and repetitions. For Figure 3i, AMBER run time was divided by GradeBins run time for each bin set and repetition; cells show medians of the three ratios. The repetitions are technical replicates, and the comparisons are descriptive.

### Ground truth from DNA and whole genes

For Figure 4a,d, source specific minimap2 2.31-r1302^58^ alignments used asm5 and explicit CIGAR output. Primary alignments with ≥99% identity (matching bases/alignment block length) supplied reference positions from M, = and X operations; deletions and skipped positions were excluded. Unique covered positions divided by source FASTA length defined DNA completeness. Separate summaries using mapping quality ≥20 or including reported alternative placements assess repeat ambiguity. Draft references and ambiguous repeat placements limit interpretation to the supplied sequence and alignment rule. Contamination is known foreign sequence length/total bin length.

Controls have exact coordinates. Native denotes sequence from the reference genome that a control represents, and foreign denotes sequence from a donor genome. DNA completeness is the number of unique retained native coordinates divided by the native reference length; contamination is foreign bases/total bin bases. Duplicated native DNA contributes no foreign DNA. For eligible annotated protein coding genes, gene completeness is the number of unique native genes fully retained divided by the original number of native genes. Gene contamination is the number of unique foreign genes fully retained divided by the number of all unique genes fully retained, and is undefined when no genes remain.

A whole gene must lie entirely within one fragment; fragments are not joined and repeated gene identities count once. Retention establishes sequence presence, not expression or function. Figure 4b,c,f,g compares the same estimates for 2,912 controls with DNA and whole gene ground truths.

### Construction of control bins

Controls used all 26 MBARC-26 strains^37^, comprising 23 bacteria and three archaea. NCBI assemblies and annotations provided 38 replicons, 104,992,012 bases and 94,107 eligible coding sequence identities; 1,470 pseudo, partial, compound or otherwise unsupported loci were excluded. Salmonella enterica RSK2980 GenBank annotations were retained and flagged because RefSeq carried an annotation completeness warning.

Before quality assessment, we fixed a design with approximately 2 or 20 kb fragments, shuffled and retained to target 25%, 50%, 75% or 100% native coverage. Foreign fragments targeted 0%, 5%, 10% or 20% of total bin DNA, using either the nearest available donor under independent taxonomy or a donor from another phylum. Two seeds yielded 2,912 bins, referred to as core controls; zero foreign DNA occurred once per retention, fragment length and seed. Because whole fragments were selected, realized ground truth can differ from the targets. Donor relationships and contributions are recorded; the fixed rule for donors from another phylum gave unequal donor representation.

Additional controls comprised 26 intact references and 312 duplication controls. The latter retained approximately 75% native DNA and added copies targeting 5%, 20% or 50% of that retained sequence, at both fragment lengths and seeds. After inspecting ground truth quantities, but before estimating quality, we fixed 728 controls near the cutoffs. These controls had 90% or 95% retention, 20 kb fragments and 0%, 2%, 5% or 8% foreign DNA, and used nearest donors and a balanced schedule of donors from other phyla.

All 3,978 FASTAs used anonymous filenames and contig identifiers, with origins stored separately. Independent reconstruction from source coordinates reproduced the sequences and counts of unique native bases, duplicates and whole genes. Unmodified BBTools 40.04 estimated all controls in 52 batch reports. Controls from the same source genome formed one resampling group.

### Comparison tools and definitions

CheckM 1.2.5 (CheckM1)^24^, CheckM2 1.1.0^25^ and GUNC 1.1.1^26^ received the same 8,595 nucleotide inputs as GradeBins (6,374 simulated bins, 173 MBARC-26 bins, 46 soil MAGs and 2,002 controls). Where seeds applied, the subset of controls used seed 4004 and comprised 26 intact references, 1,456 core controls, 156 duplication controls and 364 controls near the cutoffs. Dataset, bin set and filename identified each bin across outputs.

CheckM lineage_wf used database 2015_01_16, pplacer 1.1.alpha19^59^ and 16 workflow and placement threads. CheckM2 used its installed UniRef100^60^ database with KEGG Orthology (KO)^61^ annotations; GUNC used proGenomes 2.1^62^, DIAMOND 2.1.24^63^ and Prodigal 2.6.3^64^.

Contamination definitions were retained. CheckM1 uses excess marker copies; CheckM2 predicts contamination relative to expected complete uncontaminated genome length. GUNC contamination_portion is the fraction of retained, taxonomically assigned genes outside the largest group at the reported level, multiplied by 100 here. It differs from GUNC clade separation and pass/fail outputs. Differences from DNA contamination can reflect both the definition and estimation error.

Figure 4h compares identical bins across tools, using 6,363 simulated bins with numerical GUNC results and 728 controls per fragment length. Excluding 11 missing GUNC results retains all 1,552 source genomes. Completeness comparisons use all 6,374 simulated bins. Figure 4i,j uses 104 controls at zero foreign DNA and 208 at each nonzero level per fragment length. Figure 4k pools fragment lengths, giving 52 controls per duplication level; 46 GUNC estimates were zero and six nonzero at each level. Lines and bands show medians and interquartile ranges.

### Sequenced metagenomes

The MBARC-26 assembly^37^ and five bin sets came from the BinSPreader^38^ protocol^39^ dataset, version 3^40^. The bin sets are one MetaBAT2 bin set and four refinements of it, totalling 173 bins from one sequencing experiment. Figure 5 compares estimates without assigning bins to reference strains.

We retained all 46 deposited MAGs (151,110,167 bases) from GEM soil metagenome 3300005529^5^, selected before the new estimates because it had the most available deposited soil genomes. Original filtering used CheckM 1.0.11 completeness ≥50%, contamination <5% and completeness − 5 × contamination >50, in percentages. Figure 5 therefore compares new estimates within a previously selected collection; historical estimates remain separate metadata.

### Error summaries and uncertainty

Error is 100 × (estimated fraction − ground truth fraction), in percentage points. Mean absolute error (MAE) averages absolute errors within genomes, then weights genomes equally. Figure labels MAE (%) denote percentage points. Original estimates were retained.

Error intervals are the 2.5th–97.5th percentiles of 2,000 bootstrap samples of source genomes, retaining all bins derived from each sampled genome. A source shared across simulated communities forms one group. Contamination comparisons used identical draws across tools. Intervals are conditional on observed sources and donor design; reused donors introduce residual dependence, and annotation uncertainty is excluded. Missing values remain missing.

Figure 5a–e uses separate ordinary least squares fits with an intercept for each tool, quantity and dataset, weighting bins equally. Pointwise 95% intervals for fitted means use 2,000 bootstrap samples. MBARC-26 bins were assigned to the original MetaBAT2 bin sharing the most contig bases, with ties resolved by identifier, giving 35 groups containing all 173 bins. Each sampled group retains its bins from all five bin sets. Shared sequence across groups can leave residual dependence. Soil resampling uses 46 MAGs. Draws were shared across comparisons within datasets; all fits were valid. Lines and bands span observed GradeBins values and may exceed 0–100%. Bands describe fitted means within these collections, rather than individual predictions or replication across metagenomes.

### Quality selection and F1

The HQ screen requires completeness >90% and contamination <5%, the completeness and contamination criteria of the MIMAG high quality standard^2^. Full designation additionally requires RNA evidence. Figure 4l–n applies this screen to 364 controls with 20 kb fragments, 26 genomes and seed 4004. DNA ground truth uses unique native bases/reference length and foreign bases/total bin length. Integer checks preserve strict inequalities, identifying 167 passing and 197 failing bins. Matrix percentages use these row totals.

In Figure 4o, MAE uses 728 core controls with 20 kb fragments; selection F1 uses the separate 364 controls near the cutoffs. Passing both cutoffs defines a positive. Precision = TP/(TP+FP), recall = TP/(TP+FN) and F1 = 2TP/(2TP+FP+FN), with true positives (TP), false positives (FP) and false negatives (FN) pooled across bins. Intervals use 2,000 bootstrap samples of source genomes with shared draws across tools. F1 depends on the designed passing/failing proportions; undefined denominators remain missing. GUNC provides contamination MAE only.

Figure 5f applies HQ cutoffs to original unrounded estimates. Among bins not assigned HQ by a tool, MQ requires completeness ≥50% and contamination <10%. Classes were assigned here from the estimates of each tool with these cutoffs. Each Venn diagram counts, for bins assigned the class by at least one tool, the exact combination of tools that assigned it; circle areas are schematic. Source Data retain all 173 MBARC-26 and 46 soil decisions, including bins outside the diagrams. GUNC lacks completeness estimates and supplies no combined assignment.

### Run time and RAM measurements

Figure 6a contains 45 GradeBins runs made before the controlled comparison, one for each combination of five communities, three bin sets and three assessment modes (measurement from contig labels with the assembly or the whole genome reference, or quality estimation). Input size is total bin DNA. Hardware differed and each input/mode was run once.

The controlled comparison of quality estimation used the MetaBAT2 bin sets of 11, 111 and 1,133 simulated bins (labelled Synthetic in Fig. 6), 35 MBARC-26 bins and 46 soil MAGs. Each tool ran three times per dataset. Within each dataset and repetition, all four ran sequentially in a fixed randomized order on the same node with the same 16 logical CPUs (eight physical cores, two hardware threads each). Runs spanned seven nodes, all with AMD EPYC 9555P processors. Each job reserved all memory of its node, which excluded other jobs. GradeBins had a 16 GiB maximum Java heap; physical memory did not limit runs.

Each invocation used a fresh output directory. Timing included startup, model/database loading, gene prediction, analysis and the principal quality report, excluding installation and database preparation. Extra CheckM quality annotations and GUNC taxonomy details used in accuracy analyses were outside timed commands. Operating system caches were not flushed. All 60 runs completed.

GNU time^41^ recorded elapsed time, reported here as run time, and maximum resident set size (RSS), which on Linux is the largest recorded individual process peak, including completed child programs. Dividing KiB by 2^20^ gives GiB. Figure 6 and Extended Data Figure 2 show three runs, medians and ranges. The 16 GiB line in the figures marks the Java heap setting, which differs from total RAM.

For Figure 6d,e, comparison tool time or RAM was divided by GradeBins time or RAM for each dataset and repetition. Heatmap cells show medians of three ratios (45 ratios per quantity). GradeBins run time includes waiting for its taxonomy service. RAM covers local processes; server CPU, RAM and cache state were unmeasured.

### Combined RAM of tool and helpers

A sampler read /proc/PID/smaps_rollup and summed proportional set size (PSS) across the timed command and tracked helpers, including descendants surviving their parent. PSS apportions shared pages and excludes unmapped filesystem cache.

The sampler paused 0.5 seconds between measurements, but actual intervals reached 11.83 seconds and may miss brief peaks. Source data retain sample counts and interval summaries. Extended Data Figure 2 reports the largest sampled combined PSS and its ratio to GNU time RSS, with 15 runs per tool. Peaks can occur at different times and memory accounting differs, so ratios can be below 1. Measurements cover local processes only.

### Software compatibility and figures

GradeBins runs on controls and sequenced metagenomes, and the 15 GradeBins runs in the controlled comparison of run time and RAM, used unmodified BBTools 40.04. Two earlier estimation runs, on the two QuickBin bin sets of the planned 500 genome community, required a compatibility fix that supplied a fallback for a viral lineage without a recognized domain; with the fix, all 663 and 662 bins were retained. When the analysis of the 663-bin set was repeated, the report rows of its other 662 bins were unchanged.

Analyses used Python 3.13.12^65^ with NumPy 2.5.4^66^ and pandas 3.0.1^67,68^. Figures were drawn in R 4.5.2^69^ with tidyverse 2.0.0^70^, ggplot2 4.0.2^71^ and cowplot 1.2.0^72^.

### Running GradeBins

For internal prokaryotic estimation with an assembly and bin directory, run:

`gradebins.sh -Xmx16g threads=16 in=bins ref=assembly.fa prokcc quickclade=f report=report.tsv`

Alternatively, checkm=quality_report.tsv imports a CheckM2 report. For measurement from contig labels against whole genome lengths, taxin= supplies the table of source genome sizes and size= supplies the prepared assembly size for recovery summaries. Input and output options are documented at https://bbmap.org/tools/gradebins.

### Use of AI tools

AI assistants, including OpenAI Codex, assisted language editing, restructuring, analysis scripts, data source checks, figure preparation and organization of feedback. OpenAI ChatGPT 5.2 Pro reviewed an earlier draft. The authors are responsible for the analyses, interpretation and manuscript.

## Data availability

Supplementary Tables 1 and 2 describe simulated communities and their members. Soil genomes and historical quality metadata are from the GEM catalogue^5^, metagenome 3300005529 (https://portal.nersc.gov/GEM/). The sequenced MBARC-26 assembly and five bin sets are from the BinSPreader protocol dataset, version 3^40^ (https://doi.org/10.6084/m9.figshare.21970808.v3). Source Data tables accompany the six main and two Extended Data figures, providing observations, summaries, units, panel mappings and definitions.

## Code availability

GradeBins is distributed with BBTools (https://bbmap.org/tools/gradebins; https://github.com/bbushnell/BBTools; https://sourceforge.net/projects/bbmap/files/). We evaluated version

40.04. ProkCC source is public in the same repository (https://github.com/bbushnell/BBTools/blob/master/prokcc.sh); a separate manuscript is in preparation. The accompanying analysis archive includes scripts and notebooks, source tables, and the AMBER input files, output reports and execution records used for Figure 3.

## Author contributions

BB and JCV designed the project. BB developed GradeBins and generated the simulated data. JCV developed the benchmarking and analysis workflow, configured software on the JGI Dori cluster and drafted the manuscript. RMB contributed to the interpretation of results and contributed to the discussion. BB wrote the methods for GradeBins and simulated data generation. All authors interpreted the results and revised the manuscript.

## Correspondence

Correspondence and requests for materials should be addressed to Brian Bushnell or Juan C. Villada.

## References

1. Chen, L.-X., Anantharaman, K., Shaiber, A., Eren, A. M. & Banfield, J. F. Accurate and complete genomes from metagenomes. Genome Res. 30, 315–333 (2020).

2. Bowers, R. M. et al. Minimum information about a single amplified genome (MISAG) and a metagenome-assembled genome (MIMAG) of bacteria and archaea. Nat. Biotechnol. 35, 725–731 (2017).

3. Tyson, G. W. et al. Community structure and metabolism through reconstruction of microbial genomes from the environment. Nature 428, 37–43 (2004).

4. Parks, D. H. et al. Recovery of nearly 8,000 metagenome-assembled genomes substantially expands the tree of life. Nat Microbiol 2, 1533–1542 (2017).

5. Nayfach, S. et al. A genomic catalog of Earth’s microbiomes. Nat. Biotechnol. 39, 499–509 (2021).

6. Szabó, G., Eloe-Fadrosh, E. A., Pett-Ridge, J. & Woyke, T. A genomic view of Earth’s biomes. Nat. Rev. Genet. 27, 13–31 (2026).

7. Pasolli, E. et al. Extensive unexplored human microbiome diversity revealed by over 150,000 genomes from metagenomes spanning age, geography, and lifestyle. Cell 176, 649–662.e20 (2019).

8. Almeida, A. et al. A unified catalog of 204,938 reference genomes from the human gut microbiome. Nat. Biotechnol. 39, 105–114 (2021).

9. Shaiber, A. & Eren, A. M. Composite metagenome-assembled genomes reduce the quality of public genome repositories. mBio 10, e00725–19 (2019).

10. Li, D., Liu, C.-M., Luo, R., Sadakane, K. & Lam, T.-W. MEGAHIT: an ultra-fast single-node solution for large and complex metagenomics assembly via succinct de Bruijn graph. Bioinformatics 31, 1674–1676 (2015).

11. Nurk, S., Meleshko, D., Korobeynikov, A. & Pevzner, P. A. metaSPAdes: a new versatile metagenomic assembler. Genome Res. 27, 824–834 (2017).

12. Albertsen, M. et al. Genome sequences of rare, uncultured bacteria obtained by differential coverage binning of multiple metagenomes. Nat. Biotechnol. 31, 533–538 (2013).

13. Alneberg, J. et al. Binning metagenomic contigs by coverage and composition. Nat. Methods 11, 1144–1146 (2014).

14. Wu, Y.-W., Simmons, B. A. & Singer, S. W. MaxBin 2.0: an automated binning algorithm to recover genomes from multiple metagenomic datasets. Bioinformatics 32, 605–607 (2016).

15. Kang, D. D. et al. MetaBAT 2: an adaptive binning algorithm for robust and efficient genome reconstruction from metagenome assemblies. PeerJ 7, e7359 (2019).

16. Bushnell, B. & Villada, J. C. Deployable high-fidelity metagenome binning at scale with QuickBin. *Commun*. Biol. 9, 1135 (2026).

17. Nissen, J. N. et al. Improved metagenome binning and assembly using deep variational autoencoders. Nat. Biotechnol. 39, 555–560 (2021).

18. Líndez, P. P. et al. Adversarial and variational autoencoders improve metagenomic binning. Commun Biol 6, 1073 (2023).

19. Pan, S., Zhu, C., Zhao, X.-M. & Coelho, L. P. A deep siamese neural network improves metagenome-assembled genomes in microbiome datasets across different environments. Nat. Commun. 13, 2326 (2022).

20. Pan, S., Zhao, X.-M. & Coelho, L. P. SemiBin2: self-supervised contrastive learning leads to better MAGs for short- and long-read sequencing. Bioinformatics 39, i21–i29 (2023).

21. Wang, Z. et al. Effective binning of metagenomic contigs using contrastive multi-view representation learning. Nat. Commun. 15, 585 (2024).

22. Sieber, C. M. K. et al. Recovery of genomes from metagenomes via a dereplication, aggregation and scoring strategy. Nat Microbiol 3, 836–843 (2018).

23. Uritskiy, G. V., DiRuggiero, J. & Taylor, J. MetaWRAP-a flexible pipeline for genome-resolved metagenomic data analysis. Microbiome 6, 158 (2018).

24. Parks, D. H., Imelfort, M., Skennerton, C. T., Hugenholtz, P. & Tyson, G. W. CheckM: assessing the quality of microbial genomes recovered from isolates, single cells, and metagenomes. Genome Res. 25, 1043–1055 (2015).

25. Chklovski, A., Parks, D. H., Woodcroft, B. J. & Tyson, G. W. CheckM2: a rapid, scalable and accurate tool for assessing microbial genome quality using machine learning. Nat. Methods 20, 1203–1212 (2023).

26. Orakov, A. et al. GUNC: detection of chimerism and contamination in prokaryotic genomes. Genome Biol. 22, 178 (2021).

27. Mattock, J. & Watson, M. A comparison of single-coverage and multi-coverage metagenomic binning reveals extensive hidden contamination. Nat. Methods 20, 1170–1173 (2023).

28. Han, H., Wang, Z. & Zhu, S. Benchmarking metagenomic binning tools on real datasets across sequencing platforms and binning modes. Nat. Commun. 16, 2865 (2025).

29. Kim, J., Kim, N., Cha, J. H., Ma, J. & Lee, I. Comprehensive benchmarking of metagenomic binning tools reveals key factors for improved genome recovery. Nat. Commun. 17, 3467 (2026).

30. Sczyrba, A. et al. Critical Assessment of Metagenome Interpretation-a benchmark of metagenomics software. Nat. Methods 14, 1063–1071 (2017).

31. Meyer, F. et al. Critical Assessment of Metagenome Interpretation: the second round of challenges. Nat. Methods 19, 429–440 (2022).

32. Fritz, A. et al. CAMISIM: simulating metagenomes and microbial communities. Microbiome 7, 17 (2019).

33. Bushnell, B., Schulz, F. & Villada, J. C. RandomReadsMG: Rapid, realistic metagenome simulation at terabase scale for benchmarking and experimental design. Cell Genom. 101384 (2026).

34. Meyer, F. et al. AMBER: Assessment of Metagenome BinnERs. Gigascience 7, giy069 (2018).

35. Bushnell, B. BBTools. https://sourceforge.net/projects/bbmap/.

36. Bushnell, B. BBMap: A Fast, Accurate, Splice-Aware Aligner. https://www.osti.gov/servlets/purl/1241166 (2014).

37. Singer, E. et al. Next generation sequencing data of a defined microbial mock community. Sci. Data 3, 160081 (2016).

38. Tolstoganov, I., Kamenev, Y., Kruglikov, R., Ochkalova, S. & Korobeynikov, A. BinSPreader: Refine binning results for fuller MAG reconstruction. iScience 25, 104770 (2022).

39. Ochkalova, S., Tolstoganov, I., Lapidus, A. & Korobeynikov, A. Protocol for refining metagenomic binning with BinSPreader. STAR Protoc. 4, 102417 (2023).

40. Ochkalova, S. & Korobeynikov, A. BinSPreader binning refining protocol. figshare 10.6084/M9.FIGSHARE.21970808.V3 (2023).

41. Free Software Foundation. GNU Time. https://www.gnu.org/software/time/manual/ (2026).

42. Maguire, F. et al. Metagenome-assembled genome binning methods with short reads disproportionately fail for plasmids and genomic Islands. Microb. Genom. 6, 000436 (2020).

43. Olm, M. R., Brown, C. T., Brooks, B. & Banfield, J. F. dRep: a tool for fast and accurate genomic comparisons that enables improved genome recovery from metagenomes through de-replication. ISME J. 11, 2864–2868 (2017).

44. Bickhart, D. M. et al. Generating lineage-resolved, complete metagenome-assembled genomes from complex microbial communities. Nat. Biotechnol. 40, 711–719 (2022).

45. Sereika, M. et al. Oxford Nanopore R10.4 long-read sequencing enables the generation of near-finished bacterial genomes from pure cultures and metagenomes without short-read or reference polishing. Nat. Methods 19, 823–826 (2022).

46. Benoit, G. et al. High-quality metagenome assembly from long accurate reads with metaMDBG. Nat. Biotechnol. 42, 1378–1383 (2024).

47. Sereika, M. et al. Genome-resolved long-read sequencing expands known microbial diversity across terrestrial habitats. Nat. Microbiol. 10, 2018–2030 (2025).

48. Jin, H. et al. A high-quality genome compendium of the human gut microbiome of Inner Mongolians. Nat. Microbiol. 8, 150–161 (2023).

49. Nayfach, S., Shi, Z. J., Seshadri, R., Pollard, K. S. & Kyrpides, N. C. New insights from uncultivated genomes of the global human gut microbiome. Nature 568, 505–510 (2019).

50. Parks, D. H. et al. GTDB release 10: a complete and systematic taxonomy for 715 230 bacterial and 17 245 archaeal genomes. Nucleic Acids Res. 54, D743–D754 (2026).

51. Astashyn, A. et al. Rapid and sensitive detection of genome contamination at scale with FCS-GX. Genome Biol. 25, 60 (2024).

52. Rinke, C. et al. Insights into the phylogeny and coding potential of microbial dark matter. Nature 499, 431–437 (2013).

53. Pachiadaki, M. G. et al. Charting the complexity of the marine microbiome through single-cell genomics. Cell 179, 1623–1635.e11 (2019).

54. Saary, P., Mitchell, A. L. & Finn, R. D. Estimating the quality of eukaryotic genomes recovered from metagenomic analysis with EukCC. Genome Biol. 21, 244 (2020).

55. Chaumeil, P.-A., Mussig, A. J., Hugenholtz, P. & Parks, D. H. GTDB-Tk: a toolkit to classify genomes with the Genome Taxonomy Database. Bioinformatics 36, 1925–1927 (2020).

56. Chaumeil, P.-A., Mussig, A. J., Hugenholtz, P. & Parks, D. H. GTDB-Tk v2: memory friendly classification with the Genome Taxonomy Database. Bioinformatics 38, 5315–5316 (2022).

57. Cox, E. et al. NCBI Taxonomy: enhanced access via NCBI Datasets. Nucleic Acids Res. 53, D1711–D1715 (2025).

58. Li, H. Minimap2: pairwise alignment for nucleotide sequences. Bioinformatics 34, 3094–3100 (2018).

59. Matsen, F. A., Kodner, R. B. & Armbrust, E. V. pplacer: linear time maximum-likelihood and Bayesian phylogenetic placement of sequences onto a fixed reference tree. BMC Bioinformatics 11, 538 (2010).

60. Suzek, B. E. et al. UniRef clusters: a comprehensive and scalable alternative for improving sequence similarity searches. Bioinformatics 31, 926–932 (2015).

61. Kanehisa, M. & Goto, S. KEGG: Kyoto Encyclopedia of Genes and Genomes. Nucleic Acids Research 28, 27–30 (2000).

62. Mende, D. R. et al. proGenomes2: an improved database for accurate and consistent habitat, taxonomic and functional annotations of prokaryotic genomes. Nucleic Acids Research 48, D621–D625 (2020).

63. Buchfink, B., Reuter, K. & Drost, H.-G. Sensitive protein alignments at tree-of-life scale using DIAMOND. Nature Methods 18, 366–368 (2021).

64. Hyatt, D. et al. Prodigal: prokaryotic gene recognition and translation initiation site identification. BMC Bioinformatics 11, 119 (2010).

65. Python Software Foundation. Python 3.13.12. https://www.python.org/downloads/release/python-31312/ (2026).

66. Harris, C. R. et al. Array programming with NumPy. Nature 585, 357–362 (2020).

67. McKinney, W. Data Structures for Statistical Computing in Python. in Proceedings of the 9th Python in Science Conference (eds van der Walt, S. & Millman, J.) 56–61 (2010).

68. The pandas development team. Pandas-Dev/pandas: Pandas. (Zenodo, 2026). doi:10.5281/zenodo.18675244.

69. R Core Team. R: A Language and Environment for Statistical Computing. http://dx.doi.org/10.32614/R.manuals (2025) doi:10.32614/R.manuals.

70. Wickham, H. et al. Welcome to the tidyverse. Journal of Open Source Software 4, 1686 (2019).

71. Wickham, H. ggplot2: Elegant Graphics for Data Analysis. (Springer, Cham, 2016).

72. Wilke, C. O. Cowplot: Streamlined Plot Theme and Plot Annotations for ‘ggplot2’. https://wilkelab.org/cowplot/ (2025).

